# ALS-linked mutations impair UBQLN2 stress-induced biomolecular condensate assembly in cells

**DOI:** 10.1101/2020.10.17.335380

**Authors:** Julia F. Riley, Heidi Hehnly, Carlos A. Castañeda

## Abstract

Mutations in Ubiquilin-2 (UBQLN2), a ubiquitin-binding shuttle protein involved in several protein quality control processes, can lead to amyotrophic lateral sclerosis (ALS). We previously found that wild-type UBQLN2 forms dynamic, membraneless biomolecular condensates upon cellular stress, and undergoes liquid-liquid phase separation *in vitro*. However, the impact of ALS-linked mutations on UBQLN2 condensate formation in cells is unknown. Here, we employ live-cell imaging with photokinetic analysis to investigate how five patient-derived ALS-linked mutations in UBQLN2 impact stress-induced UBQLN2 condensate assembly and condensate material properties. Both wild-type and mutant UBQLN2 condensates are generally cytoplasmic and liquid-like. However, cells transfected with mutant UBQLN2 contain fewer stress-induced UBQLN2 condensates than those with wild-type UBQLN2. Most strikingly, ectopically expressed P506T UBQLN2 forms the lowest number of stress-induced condensates of all UBQLN2 mutants, and these condensates are significantly smaller than those of wild-type UBQLN2. Fluorescence recovery after photobleaching (FRAP) analysis of UBQLN2 condensates revealed higher immobile fractions for UBQLN2 mutants, especially P506T. P497S and P497H mutations differentially impact condensate properties, demonstrating that the effects of ALS-linked mutations are both position- and amino acid-dependent. Collectively, our data show that disease mutations hinder assembly and alter viscoelastic properties of stress-induced UBQLN2 condensates, potentially leading to aggregates commonly observed in ALS.

## Introduction

Amyotrophic lateral sclerosis (ALS) is a neurodegenerative disease distinguished by progressive, selective motor neuron death (Cleveland and Rothstein, 2001). Most (~90%) of ALS cases are classified as sporadic ALS (sALS), whereas the remaining ~10% of cases are familial ALS (fALS) cases caused by mutations in a subset of genes encoding protein quality control and RNA-binding proteins, among others (Cleveland and Rothstein, 2001; Taylor *et al*., 2016). One such fALS-linked gene encodes for the shuttle protein Ubiquilin-2 (UBQLN2) (Kleijnen *et al*., 2000; Mah *et al*., 2000). Mutations in UBQLN2, primarily in its proline-rich PXX region, underlie 1-2% of fALS cases (Deng *et al*., 2011), and cause age-related motor defects (Le *et al*., 2016; Whiteley *et al*., 2020). Although best known for its role in the ubiquitin-proteasome system (Ko *et al*., 2004; Hjerpe *et al*., 2016), UBQLN2 also participates in other protein quality control processes including endoplasmic reticulum-associated protein degradation (Lim *et al*., 2009; Xia *et al*., 2014) and macroautophagy (Chen *et al*., 2018; Wu *et al*., 2020).

ALS-linked UBQLN2 mutations have been shown to perturb both proteasomal degradation and autophagy processes. One mutation, P497H, causes accumulation of autophagy cargo protein p62 and autophagosome marker LC3-II (Wu *et al*., 2015). Other mutations, including P497S, are deficient in binding heat shock protein 70 (HSP-70) (Teyssou *et al*., 2017), and therefore may affect the ability of UBQLN2 to clear protein aggregates (Hjerpe *et al*., 2016). Wild-type (WT) UBQLN2 is also implicated as a participant in ALS pathology because it may interact directly with TDP-43, a protein whose aggregation is associated with > 95% of ALS cases, and promote TDP-43 assembly into cytoplasmic inclusions (Deng *et al*., 2011; Cassel and Reitz, 2013). It remains unclear whether mutation-induced functional deficiencies converge on any singular, critical function.

Endogenously expressed UBQLN2 colocalizes with stress granules, whereas ectopically expressed UBQLN2 also forms stress-induced biomolecular condensates that do not contain stress granule markers (Alexander *et al*., 2018; Dao *et al*., 2018). Stress granules are dynamic ribonucleoprotein condensates that form in response to external stressors such as arsenite-induced oxidative stress, osmotic stress, and heat shock (Buchan and Parker, 2009). Importantly, ALS-linked proteins such as TDP-43, FUS, TIA-1, and hnRNPA1, many of which interact with UBQLN2, also form biomolecular condensates, are recruited to stress granules, and localize to protein aggregates (Molliex *et al*., 2015; Patel *et al*., 2015; Lee *et al*., 2016; Boeynaems *et al*., 2017; Mackenzie *et al*., 2017; Gasset-Rosa *et al*., 2019; Mann *et al*., 2019). Together, these findings suggest a recurring, intimate link between condensate assembly and ALS pathology.

The impact of different ALS-linked mutations in UBQLN2 on the assembly of stress-induced biomolecular condensates remains unclear. Furthermore, the dynamics of UBQLN2-containing biomolecular condensates have not been thoroughly characterized. Here, we use a live-cell imaging approach to analyze the impact of five patient-derived ALS-linked mutations in several distinct UBQLN2 domains (specifically A282V, M446R, P497H, P497S and P506T) on the assembly and material properties of UBQLN2 biomolecular condensates in cells. We found that while all UBQLN2 variants formed liquid-like biomolecular condensates in U2OS cells, mutant UBQLN2 (particularly P506T) formed fewer condensates per cell than wild-type. Fluorescence recovery after photobleaching (FRAP) analysis suggests that many mutations, most notably P506T, decreased liquidity of UBQLN2 condensates. The types of mutations also matter, as P497H and P497S mutations had different effects on condensate properties. Together, these data implicate a hindered capacity for UBQLN2 to undergo biomolecular condensate formation as a potential pathogenic mechanism in ALS.

## Materials and Methods

### Recombinant DNA

Patient-derived ALS-linked mutations (A282V, M446R, P497H, P497S, and P506T) were generated from UBQLN2 in a pmCherry-C1 vector using a Phusion Site-Directed Mutagenesis Kit (Thermo Scientific). Transfection-grade plasmid DNA were isolated using a PureLink Maxiprep Kit (ThermoFisher).

### Cell Culture and Stress Induction Experiments

A U2OS cell line (kindly provided by the J. Paul Taylor lab, St. Jude Children’s Research Hospital) stably expressing G3BP1-GFP, a fluorescently tagged stress granule marker, was used for all experiments. Cells were grown in DMEM high glucose media (ThermoFisher) supplemented with 10% Seradigm FBS (VWR). Cells were maintained at 37°C and 5% CO_2_. Twenty-four hours after plating on glass-bottom dishes (MatTek Corporation), cells were transfected. To do so, mCherry-UBQLN2 (mCh-UBQLN2) plasmids were diluted in Opti-MEM Reduced Serum Medium (ThermoFisher) and allowed 20 minutes to form complexes with Trans-IT transfection reagent (Mirus). To induce oxidative stress, sodium (meta)arsenite (NaAsO_2_, Sigma-Aldrich) was diluted in Leibovitz’s L-15 medium (ThermoFisher) supplemented with 10% FBS and filtered with a 0.2 μm filter. Cells were imaged in L-15 media supplemented with 10% FBS prior to and following the application of 0.5 mM sodium arsenite stress.

### Live-Cell Imaging

Live images of cells under stress and FRAP experiments (described below) were acquired using a Leica DMi8 STP800 (Leica, Bannockburn, IL) equipped with a X-light V2 Confocal Unit spinning disk spinning disk unit. The microscope is equipped with a Lumencor SPECTRA X (Lumencor, Beaverton, OR) with a Hamamatsu ORCAflash 4.0 V2 CMOS C11440-22CU camera and an 89 North–LDI laser with a Photometrics Prime-95B camera. The optics equipment used was an HC PL APO 63×/1.40 NA oil CS2 oil emersion objective.

Z-stacks with a 1 μm step size covering a range of 4 μm were acquired at each timepoint. Images of UBQLN2 membraneless body formation were acquired with one-minute intervals between Z-stacks. Live-cell image acquisition of the formation of mCherry-UBQLN2 (mCh-UBQLN2) bodies began directly after the addition of 0.5 mM arsenite stress.

### Image Analysis

Live images were analyzed and are presented as average intensity projections of the Z-stack (1 μm steps, 4 μm range, 5 steps total due to an additional image acquired in the center of the range). Analysis of live images was conducted using the threshold and particle analysis features on Fiji (version 2.0.0) (Schindelin *et al*., 2012). Images were converted to 8-bit, and threshold was adjusted appropriately on an individual basis to minimize the detection of background fluorescence, thus including puncta and excluding diffuse, cytosolic mCh-UBQLN2. To analyze the number of puncta and timing of their formation, one to three cells from each trial that did not contain puncta prior to arsenite-induced stress were analyzed for each mCh-UBQLN2 construct.

Fiji was used to analyze individual UBQLN2 body area, circularity and intensity from 16-bit images at 60 minutes following the application of 0.5mM arsenite. To do so, a selection was created around the three largest UBQLN2 bodies in each cell. The measure tool on Fiji was then used to determine mean intensity and area. Integrated intensity was determined by multiplying mean intensity and area measurements.

### Fluorescence Recovery After Photobleaching (FRAP) Experiments

Fluorescence recovery after photobleaching (FRAP) experiments were conducted 48-72 hours following transfection using the Leica DMi8 STP800 described above. FRAP was conducted between one hour and fifteen minutes and one hour and forty-five minutes following the addition of 0.5mM arsenite. Photobleaching with a 405 nm laser was executed using two iterations separated by 15 ms. Excitation laser exposure time for live image acquisition was 10 ms. Live image frames of the time series were acquired at 15 ms intervals.

To analyze FRAP experiments, the fluorescence intensity within the photobleached region (in this case, whole mCh-UBQLN2 puncta) was measured for each frame using 16-bit images. Five puncta were measured per cell, and five cells containing mCh-UBQLN2 puncta were assessed per trial for a total of 25 puncta per trial (n=2 trials). Normalized fluorescence recovery values for each timepoint were calculated using the equation [(*F_t_* – *F*_0_)/(*F_i_* – *F*_0_)] where F_t_ is mean fluorescence intensity at timepoint t, F_0_ is mean fluorescence intensity directly after photobleaching, and F_i_ is initial mean fluorescence intensity before photobleaching. Normalized fluorescence recovery curves of individual puncta were fitted to the one-phase association exponential model using GraphPad Prism (Version 8.3.0). Based on the fit, halftimes of fluorescence recovery (T_1/2_) were recorded, and immobile fractions were calculated by subtracting the fitted fluorescence plateau value from one.

## Results and Discussion

### WT UBQLN2 assembles into stress-induced cytoplasmic biomolecular condensates

We ectopically expressed mCherry-tagged UBQLN2 (mCh-UBQLN2; Figure 1A) in U2OS cells stably expressing G3BP-GFP, a stress granule marker. These cells enable live-imaging of stress granule formation (Yang *et al*., 2020). We subjected transfected U2OS cells to 0.5 mM arsenite stress, and simultaneously observed formation of separate G3BP-GFP and mCh-UBQLN2 puncta over the course of 60+ minutes. Whereas G3BP-marked stress granules formed quickly (10-15 minutes post stress), mCh-UBQLN2 reorganized into distinct cytoplasmic, round puncta (hereafter called UBQLN2 bodies) more slowly, approximately 30 minutes post stress (Figure 1A). UBQLN2 bodies exhibit liquid-like properties as they fuse with each other over time to form larger, rounded puncta (Figure 1B). Consequently, we classified these UBQLN2 bodies as biomolecular condensates. Our observations agree with our previous *in vitro* studies that show UBQLN2 can undergo phase separation, the biophysical process that underlies formation of biomolecular condensates (Dao *et al*., 2018).

**Figure 1.**
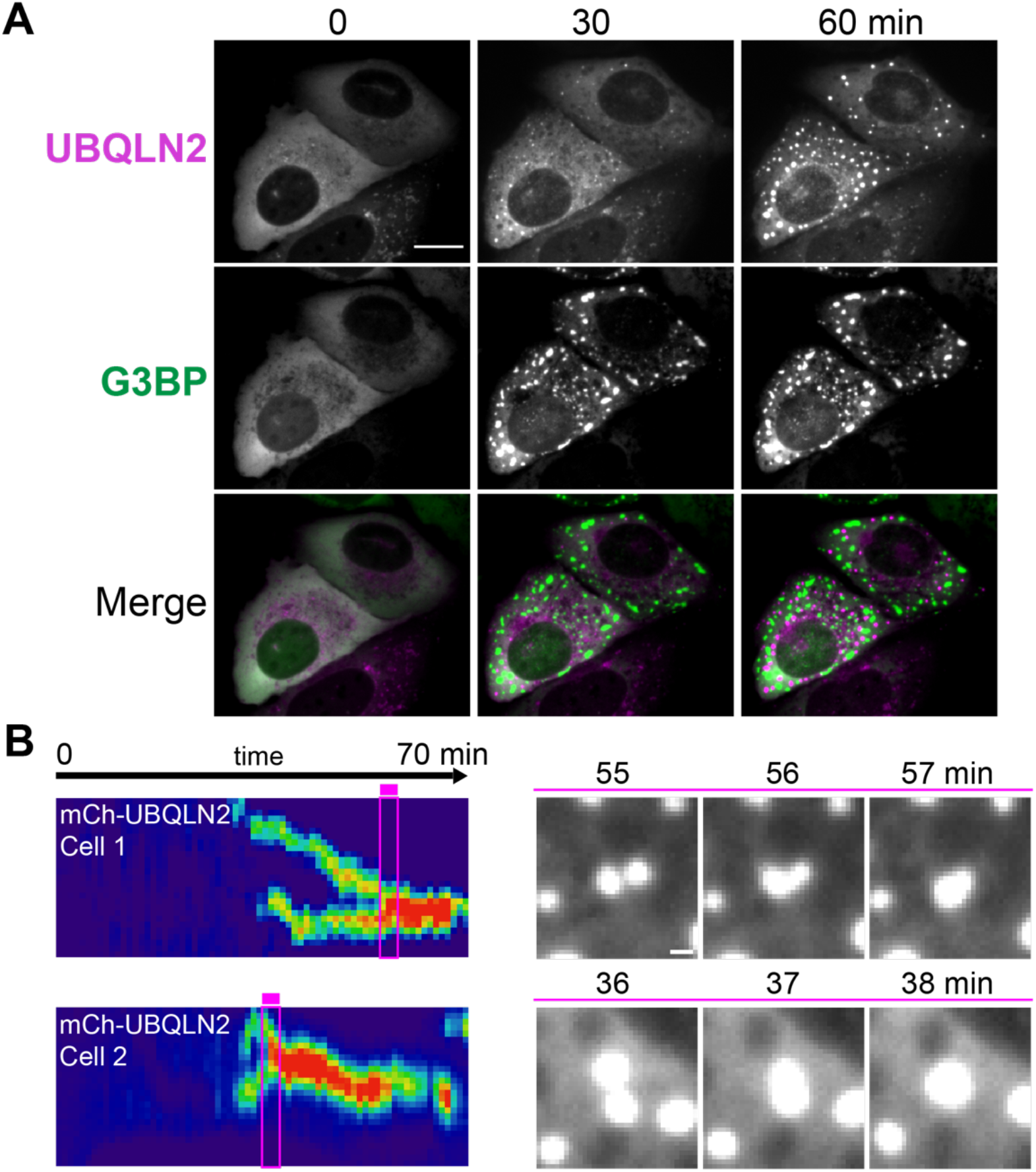
WT mCh-UBQLN2 forms liquid-like membraneless bodies in response to stress. (A) mCh-UBQLN2 and G3BP (stress granule marker) 0, 30, and 60 minutes following the addition of 0.5mM NaAsO_2_ stress. Scale bar 15 μm. (B) Representative kymographs and images of WT mCh-UBQLN2 puncta fusing; pink boxes correspond to the representative droplet fusion images shown on right. Scale bar 1 μm.

We note that mCh-UBQLN2 bodies did not colocalize with G3BP-marked stress granules; in some cases, these UBQLN2 bodies are positioned adjacent to stress granules (Figure S1A-C). These results are consistent with our prior observations for exogenously-introduced mCh-UBQLN2 in HeLa cells (Dao *et al*., 2018). Transient expression of other fluorescently-tagged phase-separating proteins or coexpression with a binding partner can sometimes result in localization to condensates distinct from those occupied by endogenously expressed proteins (Marzahn *et al*., 2016; Morelli *et al*., 2017; Bouchard *et al*., 2018). These condensates exhibit similar liquid-like properties as their endogenous counterparts and their formations are hindered by loss-of-function disease mutations. Therefore, we believe that probing the formation and properties of UBQLN2 bodies can enable an understanding of how different factors, such as stress and mutations, affect UBQLN2 in cells.

Transient expression of mCh-UBQLN2 led to two populations of cells prior to the addition of arsenite stress: cells with diffuse mCh-UBQLN2 signal throughout the cytoplasm and others with pre-existing cytoplasmic mCh-UBQLN2 puncta (Figure S1D). When transfecting a mCherry-only control, we did not observe many mCh-puncta, suggesting that puncta formation is predominantly attributed to the behavior of UBQLN2, consistent with prior studies that report on UBQLN2 puncta (Deng *et al*., 2011; Picher-Martel *et al*., 2015; Hjerpe *et al*., 2016; Sharkey *et al*., 2018). To characterize the stress-induced assembly of UBQLN2 bodies, we focused our studies on cells that exhibited diffuse mCh-UBQLN2 prior to addition of arsenite.

### Mutant UBQLN2 form fewer stress-induced condensates than WT UBQLN2

We examined the role of UBQLN2 disease-linked point mutations (A282V, M446R, P497H, P497S and P506T) on the ability of mCh-UBQLN2 to assemble oxidative stress-induced condensates following 0.5mM arsenite exposure. We chose these representative mutations because they are scattered throughout several distinct UBQLN2 domains (Figure 2A). For all mutants except P506T, we observed the formation of stress-induced UBQLN2 bodies with liquid-like and round characteristics (Figure 2B, Movies S1, S2). Cytoplasmic UBQLN2 bodies readily fused with each other to form larger, round puncta or progressively grew in size (Movies S1, S2). Strikingly, we observed very few stress-induced UBQLN2 bodies for P506T (see below).

**Figure 2.**
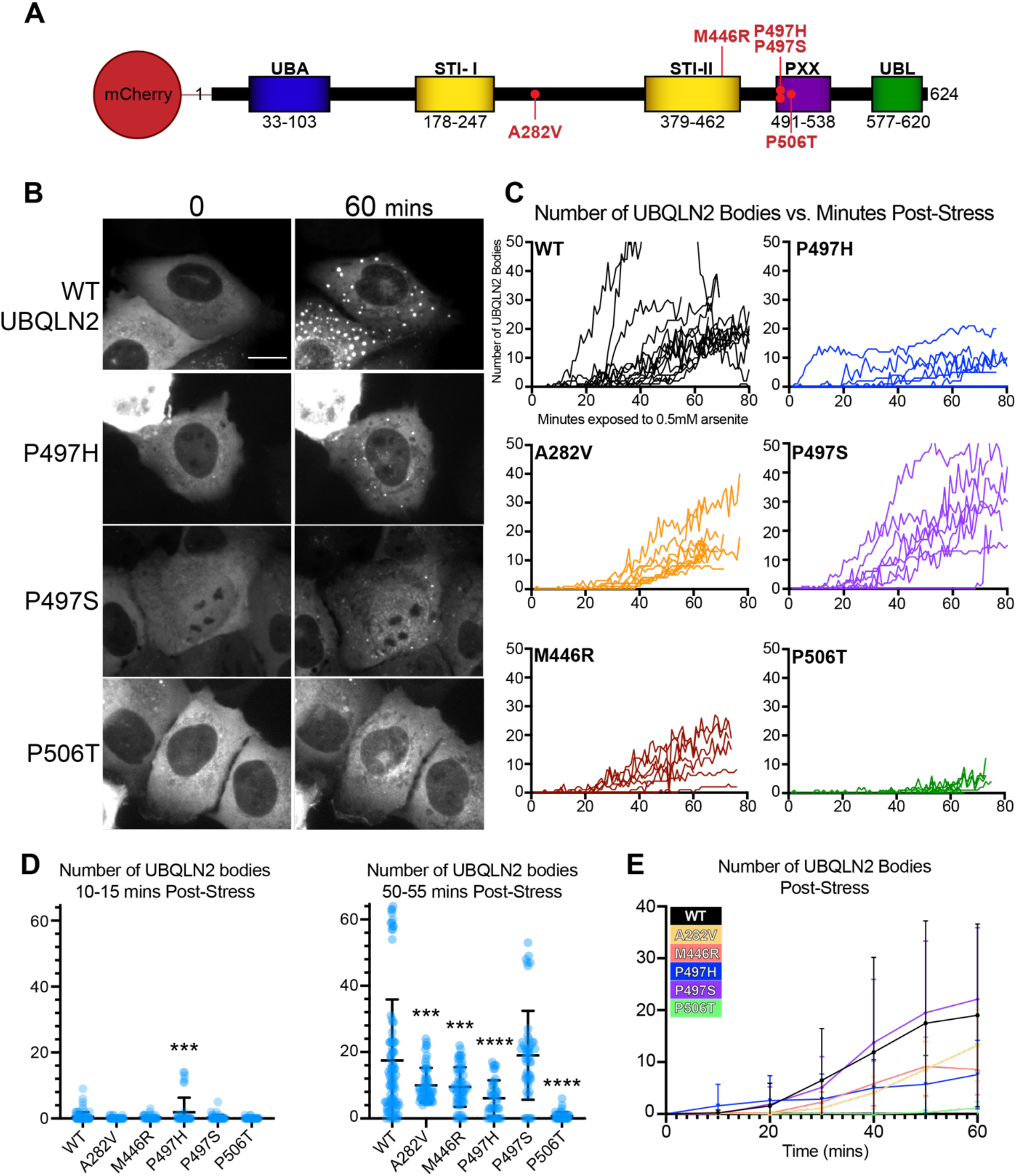
ALS-linked mutations in mCh-UBQLN2 differentially impact ability to form UBQLN2 bodies in response to stress. (A) Domain architecture of mCh-UBQLN2, including each of the five ALS-linked mutations investigated. (B) Representative images of cells before and after 60 minutes of exposure to 0.5mM arsenite. Scale bar 15 μm. (C) Number of UBQLN2 bodies formed within individual cells by WT and mutant UBQLN2 constructs after exposure to 0.5mM arsenite (WT, n= 15; A282V, n= 10; M446R, n=7; P497H, n=7; P497S, n=8; P506T, n=8). (D) Number of UBQLN2 bodies per cell between 10-15 minutes and 50-55 minutes following arsenite stress. (E) Average number of UBQLN2 bodies per cell at 10-minute intervals following the application of arsenite stress. Significance determined using one-way ANOVA with multiple comparisons to WT; * p<0.05, ** p<0.01, *** p<0.001, **** p<0.0001. Error shown as +/- SD.

To determine whether disease-linked mutations affected the stress-induced assembly of UBQLN2 bodies, we monitored both the timing of appearance for UBQLN2 bodies, as well as the number of UBQLN2 bodies per cell (Figure 2C). For WT and most of the mutants, we generally saw the onset of UBQLN2 body formation at about 30 minutes post stress. However, very few UBQLN2 bodies were observed for P506T even after 60 minutes post stress. To control for effects of UBQLN2 overexpression on possible condensate assembly, we compared the number of UBQLN2 bodies formed and the expression level per cell, but found no or weak correlation (Figure S2).

When analyzing the number of UBQLN2 bodies formed, we observed that four (A282V, M446R, P497H, and P506T) of the five mutants had significantly fewer UBQLN2 bodies per cell than WT after 50-55 minutes of exposure to arsenite stress (Figure 2D, Figure 2E). This trend is also visible on a cell-to-cell basis over time (Figure 2C). To determine whether ALS-linked mutations impact UBQLN2 condensate composition, we quantified condensate size and relative amounts of UBQLN2 within condensates by measuring fluorescence intensity post-arsenite treatment (Figure 3). While the number of stress-induced UBQLN2 bodies decreased for A282V, M446R, and P497H mutants (Figure 2D), there was little or no difference in the size of these condensates compared to WT (Figure 3B). Moreover, there was no significant difference in fluorescence intensity, indicating that the concentration of WT and mutant UBQLN2 protein are similar in these bodies. These results suggest that the decrease in the number of total stress-induced UBQLN2 bodies is not compensated by increasing the size of the UBQLN2 condensates or concentration of UBQLN2 molecules.

**Figure 3.**
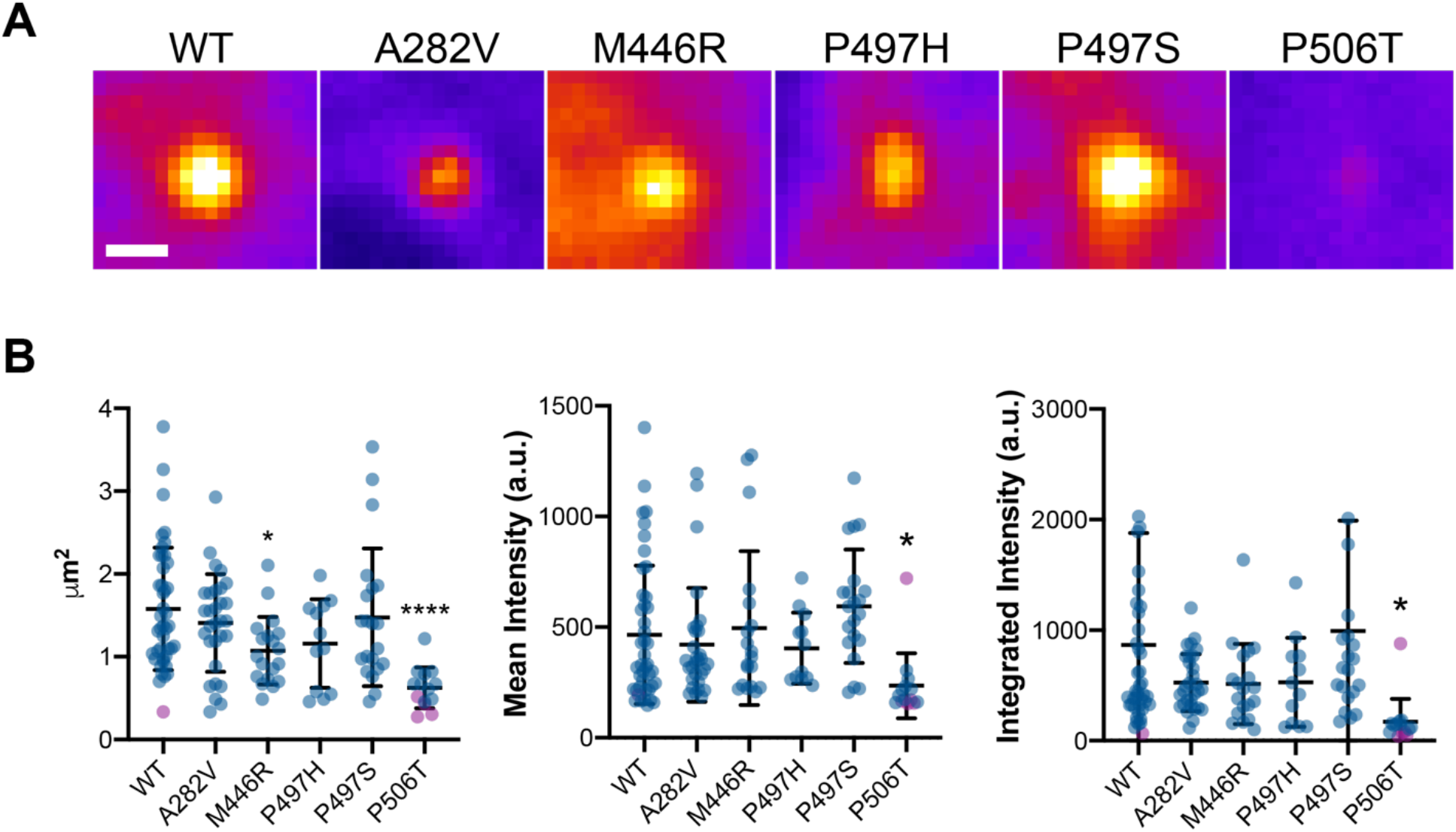
Characteristics of stress-induced, individual UBQLN2 bodies. (A) Representative images of UBQLN2 bodies 60 minutes following application of 0.5mM arsenite stress using fire LUT. Scale bar 1 μm. (B) Area, mean intensity and integrated intensity of the largest three puncta from each cell measured at 60 minutes post 0.5mM arsenite. Purple points indicate UBQLN2 bodies in cells containing less than three UBQLN2 bodies (WT, n=43; A282V, n=30; M446R, n=19, P497H, n=12; P497S, n=21; P506T, n=14). Significance determined using oneway ANOVA with multiple comparisons to WT; * p<0.05, ** p<0.01, *** p<0.001, **** p<0.0001. Error shown as +/- SD.

### P506T mutation in UBQLN2 reduces the number, size, and intensity of UBQLN2 condensates

Across our experiments, we consistently observed that the P506T mutation produced very few stress-induced mCh-UBQLN2 bodies in cells (Figure 2). Furthermore, of all mutants, P506T bodies that did form were significantly smaller and exhibited low fluorescent intensity compared to WT UBQLN2 (Figure 3). These data suggest that stress-induced condensate assembly may be impaired with the P506T mutation in UBQLN2. P506T UBQLN2 has an increased propensity to form puncta in neurons and aggregate into fibrils compared to WT UBQLN2 (Sharkey *et al*., 2018). In neurons and mammalian cell culture, P506T puncta also were notably less dynamic, larger, and more irregularly shaped than WT UBQLN2 puncta (Sharkey *et al*., 2018). This research suggests that P506T UBQLN2 is already predisposed to form puncta in cells. Thus, the P506T mutation may hinder UBQLN2’s ability to dynamically form condensates in response to external stressors. We speculate that the inability of P506T UBQLN2 to assemble into condensates under stress confers a loss of function to UBQLN2 that can lead to disease states. Indeed, P506T is one of the most deleterious UBQLN2 mutations with an early mean age of disease onset in humans (Deng *et al*., 2011; Higgins *et al*., 2019), while P506T UBQLN2 transgenic mouse models often display severe motor deficits (Le *et al*., 2016). Our data suggest that other ALS-linked mutations affect condensate behavior but to lesser degrees than P506T.

### Amino acid substitutions at position 497 affect UBQLN2 condensates differently

We observed that both the number and formation of P497S UBQLN2 bodies post-stress were strikingly similar to those of WT, suggesting that the P497S mutation did not affect the formation of stress-induced UBQLN2 bodies (Figure 2C). However, P497S additionally formed nuclear stress-induced UBQLN2 puncta, which was not observed for WT and other mutants (Movie S1, S2). Furthermore, we note differences in the overall number of stress-induced UBQLN2 bodies for P497H and P497S (Figure 2C). Therefore, amino acid substitutions at the same position (P497) differentially affect how mCh-UBQLN2 bodies form. These results are consistent with other studies that show disparate effects of single amino acid substitutions on UBQLN2 aggregation propensity (Kim *et al*., 2018; Dao *et al*., 2019) and of disease-linked mutations on aggregation and condensate behavior of other proteins such as hnRNPA1 and TDP-43 (Molliex *et al*., 2015; Conicella *et al*., 2016).

### ALS-linked mutations decrease liquid-like characteristics of mCh-UBQLN2 bodies

To assess the liquidity of UBQLN2, fluorescence recovery after photobleaching (FRAP) experiments were performed after subjecting cells to 60 minutes of arsenite stress (Figure 4). We found that the majority of WT mCh-UBQLN2 puncta (82.4%) recovered with consistently rapid kinetics (T_1/2_ = 0.8 sec) and a mobile fraction greater than 0.4 (Figure 4B, 4C). A small fraction of WT mCh-UBQLN2 condensates did not recover at all (Figure 4C). While we imaged cells after stress induction, it is possible that those UBQLN2 puncta with little fluorescent recovery may not be stress-induced puncta, but rather other UBQLN2-enriched structures in cells. Indeed, UBQLN2 is known to colocalize with other cellular structures such as autophagosomes and the ER (Wu *et al*., 2020).

**Figure 4.**
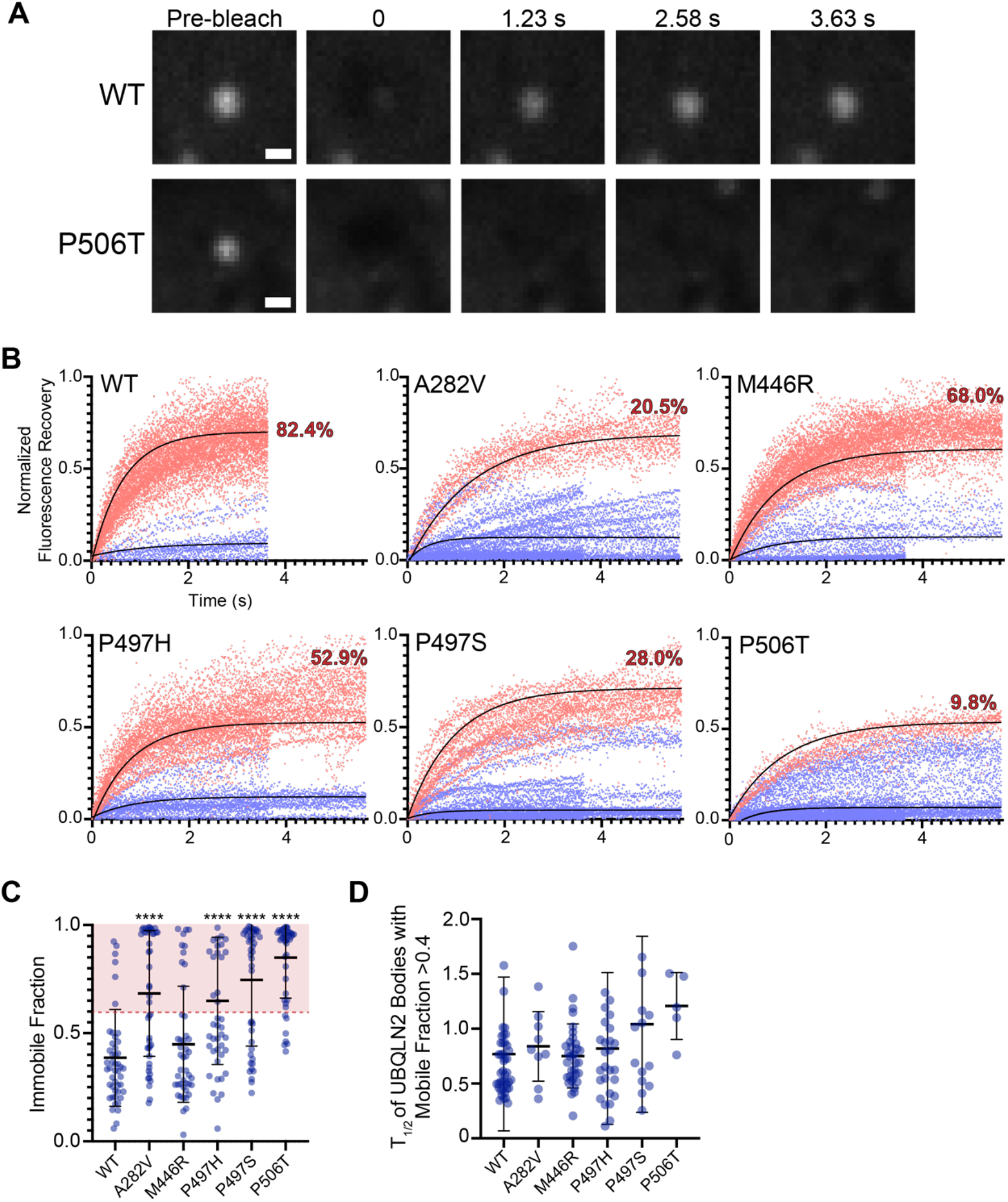
FRAP analysis of WT and mutant mCh-UBQLN2 bodies. (A) Representative images of WT and P506T mCh-UBQLN2 photobleaching experiments. Scale bar 1 μm. (B) Normalized fluorescence recovery measurements for individual UBQLN2 bodies with a recovery (mobile) fraction >0.4 (red) and <0.4 (blue), and the percentage of UBQLN2 bodies with a final mobile fraction >0.4. WT, n=51; A282V, n=44; M446R, n=50; P497H, n=51; P497S, n=50; P506T, n=50. For each experiment (n=2), a total of ~25 puncta were photobleached from five different cells. (C) Immobile fractions of UBQLN2 bodies calculated from one-phase association lines of best fit. WT, n=48; A282V, n=43; M446R, n=45; P497H, n=48; P497S, n=50; P506T, n=47. (D) Half-time of fluorescence recovery for individual UBQLN2 bodies with a mobile fraction >0.4. WT, n=42; A282V, n=9; M446R, n=34; P497H, n=27; P497S, n=14; P506T, n=5. Significance determined using one-way ANOVA with multiple comparisons to WT; * p<0.05, ** p<0.01, *** p<0.001, **** p<0.0001. Error shown as +/- SD.

A282V, P497H, P497S and P506T mCh-UBQLN2 puncta exhibited significantly higher immobile fractions than WT (Figure 4C). There was a substantial change in the distribution of UBQLN2 puncta that were classified as “recovering” (immobile fraction < 0.6) or “non-recovering” (immobile fraction > 0.6). Compared to only 17.6% non-recovering WT mCh-UBQLN2 condensates, we observed 79.5%, 47.1%, 72% and 90.2% non-recovering A282V, P497H, P497S and P506T mCh-UBQLN2 puncta, respectively (Figure 4B, 4C). We observed that P506T UBQLN2 puncta exhibited the highest immobile fraction, indicating that UBQLN2 molecules inside these puncta were the least able to exchange with surroundings, perhaps due to increased viscosity or propensity for aggregation. Our data indicate that P506T mutation affects both assembly of stress-induced UBQLN2 bodies and material properties of UBQLN2 condensates, whereas the A282V, P497H, and P497S mutations affect stress-induced condensate properties to an intermediate extent. Consistent with our observations, the P497H, P497S and P506T mutants of UBQLN2 all exhibit increased propensity to aggregate and/or form inclusions (Le *et al*., 2016; Kim *et al*., 2018).

We further examined the properties of recovering mutant UBQLN2 bodies with immobile fraction < 0.6 (Figure 4C). We observed that the fluorescence recovery half-times (T_1/2_) were similar across all mutants and WT with values ranging between 0.8 and 1.2 seconds (Figure 4D), suggesting similar kinetics for any exchange that did occur. Of the five mutants studied, the M446R variant was the only one with FRAP characteristics indistinguishable from WT (Figure 4C, 4D). M446R is a rare UBQLN2 mutation, and information about its relationship to ALS pathology is limited (Gellera *et al*., 2013). The data acquired by FRAP are indicative (although not determinant) of viscoelastic properties of membraneless bodies. Our data suggest that ALS-linked mutations alter material properties of stress-induced UBQLN2 bodies in cells. However, there remains a subset of mutant UBQLN2 bodies that exhibit WT-like fluorescence photobleaching kinetics. The impaired ability of UBQLN2 to cross between the inside and outside of puncta may affect its cellular functions, contributing to disease mechanisms.

### Conclusions

Overall, our data show that ALS-linked mutations in UBQLN2 disrupt the assembly and material properties of UBQLN2 condensates. This effect is position- and amino-acid dependent. ALS-linked mutations in UBQLN2 disrupt interactions with the proteasome, HSP70, and RNA-binding proteins, each of which is typically found in stress granules and other stress-induced condensates in cells (Chang and Monteiro, 2015; Gilpin *et al*., 2015; Hjerpe *et al*., 2016). We speculate that mutations in UBQLN2 alter these and other protein-protein interactions inside stress-induced UBQLN2 bodies, thus affecting condensate assembly and material properties. Indeed, ALS-linked mutations P497S and P506T interfered with UBQLN2’s ability to increase dynamics of FUS-RNA complexes and dysregulated stress granule formation (Alexander *et al*., 2018). Furthermore, post-translational modifications may alter UBQLN2’s ability to form condensates, similar to how phosphorylation of FUS alter condensate assembly in cells (Monahan *et al*., 2017).

We showed that ALS-linked mutant UBQLN2 condensates do not rapidly exchange contents with their surroundings, unlike WT UBQLN2 condensates. Although some ALS-linked UBQLN2 mutants have been previously demonstrated to be aggregate-prone (Kim *et al*., 2018; Sharkey *et al*., 2018; Dao *et al*., 2019), they may also be toxic in that the mutants may be less able to dynamically respond to external stressors to assemble into liquid-like condensates than WT UBQLN2. Malfunctioning UBQLN2 may lead to increased susceptibility to toxicity. UBQLN2 aggregation may be the long-term impact, and the precedent could be an alteration in adaptivity of condensate assembly to external cues. Indeed, the dynamic nature of phase separation underlies its utility in cellular stress response and potentially confers cytoprotective effects. We propose that a major function of UBQLN2 is to dynamically assemble into biomolecular condensates in response to cellular conditions.

## Author Contributions

J.F.R. and C.A.C. conceived the studies. J.F.R, H.H., and C.A.C. designed experiments. J.F.R. performed all experiments and data analysis with contributions from other authors. J.F.R. and C.A.C. wrote the paper with edits from H.H..

## Acknowledgements

We gratefully acknowledge Dr. J.P. Taylor at St. Jude’s Children’s Research Hospital for kindly providing the G3BP1-GFP U2OS cell line, and Dr. S.J. Hewett for providing laboratory space to conduct cell culture work. We thank Dr. Thuy P. Dao for assistance in generating mutant mCh-UBQLN2 constructs. This work was supported by the Beverly Petterson Bishop Neuroscience Fellowship and the Renée Crown University Honors Program Lynne Parker Funding Award to J.F.R.. This work was further supported by ALS Association grants 17-IIP-369, 18-IIP-400, and NIH R01GM136946 to C.A.C. and NIH grants R00GM107355, R01GM127621 to H.H.. This work was supported by the U.S. Army Medical Research Acquisition Activity through the FY16 Prostate Cancer Research Programs under Award W81XWH-17-1-0241 to H.H..

## References

Alexander, EJ, Niaki, AG, Zhang, T, Sarkar, J, Liu, Y, Nirujogi, RS, Pandey, A, Myong, S, and Wang, J (2018). Ubiquilin 2 modulates ALS/FTD-linked FUS–RNA complex dynamics and stress granule formation. Proc Natl Acad Sci 115, E11485–E11494.

Boeynaems, S et al. (2017). Phase Separation of C9orf72 Dipeptide Repeats Perturbs Stress Granule Dynamics. Mol Cell 65, 1044–1055.e5.

Bouchard, JJ et al. (2018). Cancer Mutations of the Tumor Suppressor SPOP Disrupt the Formation of Active, Phase-Separated Compartments. Mol Cell 72, 19–36.e8.

Buchan, JR, and Parker, R (2009). Eukaryotic Stress Granules: The Ins and Outs of Translation. Mol Cell 36, 932–941.

Cassel, JA, and Reitz, AB (2013). Ubiquilin-2 (UBQLN2) binds with high affinity to the C-terminal region of TDP-43 and modulates TDP-43 levels in H4 cells: characterization of inhibition by nucleic acids and 4-aminoquinolines. Biochim Biophys Acta 1834, 964–971.

Chang, L, and Monteiro, MJ (2015). Defective Proteasome Delivery of Polyubiquitinated Proteins by Ubiquilin-2 Proteins Containing ALS Mutations. PLoS ONE 10, e0130162.

Chen, T, Huang, B, Shi, X, Gao, L, and Huang, C (2018). Mutant UBQLN2P497H in motor neurons leads to ALS-like phenotypes and defective autophagy in rats. Acta Neuropathol Commun 6, 122.

Cleveland, DW, and Rothstein, JD (2001). From Charcot to Lou Gehrig: deciphering selective motor neuron death in ALS. Nat Rev Neurosci 2, 806–819.

Conicella, AE, Zerze, GH, Mittal, J, and Fawzi, NL (2016). ALS Mutations Disrupt Phase Separation Mediated by α-Helical Structure in the TDP-43 Low-Complexity C-Terminal Domain. Structure 24, 1537–1549.

Dao, TP, Kolaitis, R-M, Kim, HJ, O’Donovan, K, Martyniak, B, Colicino, E, Hehnly, H, Taylor, JP, and Castañeda, CA (2018). Ubiquitin Modulates Liquid-Liquid Phase Separation of UBQLN2 via Disruption of Multivalent Interactions. Mol Cell 69, 965–978.e6.

Dao, TP, Martyniak, B, Canning, AJ, Lei, Y, Colicino, EG, Cosgrove, MS, Hehnly, H, and Castañeda, CA (2019). ALS-Linked Mutations Affect UBQLN2 Oligomerization and Phase Separation in a Position- and Amino Acid-Dependent Manner. Structure 27, 937–951.

Deng, H-X et al. (2011). Mutations in UBQLN2 cause dominant X-linked juvenile and adult-onset ALS and ALS/dementia. Nature 477, 211–215.

Fecto, F, and Siddique, T (2012). UBQLN2/P62 cellular recycling pathways in amyotrophic lateral sclerosis and frontotemporal dementia. Muscle Nerve 45, 157–162.

Gasset-Rosa, F, Lu, S, Yu, H, Chen, C, Melamed, Z, Guo, L, Shorter, J, Da Cruz, S, and Cleveland, DW (2019). Cytoplasmic TDP-43 De-mixing Independent of Stress Granules Drives Inhibition of Nuclear Import, Loss of Nuclear TDP-43, and Cell Death. Neuron 102, 339–357.e7.

Gellera, C et al. (2013). Ubiquilin 2 mutations in Italian patients with amyotrophic lateral sclerosis and frontotemporal dementia. J Neurol Neurosurg Psychiatry 84, 183–187.

Gilpin, KM, Chang, L, and Monteiro, MJ (2015). ALS-linked mutations in ubiquilin-2 or hnRNPA1 reduce interaction between ubiquilin-2 and hnRNPA1. Hum Mol Genet 24, 2565–2577.

Higgins, N, Lin, B, and Monteiro, MJ (2019). Lou Gehrig’s Disease (ALS): UBQLN2 Mutations Strike Out of Phase. Structure 27, 879–881.

Hjerpe, R et al. (2016). UBQLN2 Mediates Autophagy-Independent Protein Aggregate Clearance by the Proteasome. Cell 166, 935–949.

Kim, SH, Stiles, SG, Feichtmeier, JM, Ramesh, N, Zhan, L, Scalf, MA, Smith, LM, Bhan Pandey, U, and Tibbetts, RS (2018). Mutation-dependent aggregation and toxicity in a Drosophila model for UBQLN2-associated ALS. Hum Mol Genet 27, 322–337.

Kleijnen, MF, Shih, AH, Zhou, P, Kumar, S, Soccio, RE, Kedersha, NL, Gill, G, and Howley, PM (2000). The hPLIC Proteins May Provide a Link between the Ubiquitination Machinery and the Proteasome. Mol Cell 6, 409–419.

Ko, HS, Uehara, T, Tsuruma, K, and Nomura, Y (2004). Ubiquilin interacts with ubiquitylated proteins and proteasome through its ubiquitin-associated and ubiquitin-like domains. FEBS Lett 566, 110–114.

Le, NTT et al. (2016). Motor neuron disease, TDP-43 pathology, and memory deficits in mice expressing ALS–FTD-linked UBQLN2 mutations. Proc Natl Acad Sci 113, E7580–E7589.

Lee, K-H et al. (2016). C9orf72 Dipeptide Repeats Impair the Assembly, Dynamics, and Function of Membrane-Less Organelles. Cell 167, 774–788.e17.

Lim, PJ et al. (2009). Ubiquilin and p97/VCP bind erasin, forming a complex involved in ERAD. J Cell Biol 187, 201–217.

Mackenzie, IR et al. (2017). TIA1 Mutations in Amyotrophic Lateral Sclerosis and Frontotemporal Dementia Promote Phase Separation and Alter Stress Granule Dynamics. Neuron 95, 808–816.e9.

Mah, AL, Perry, G, Smith, MA, and Monteiro, MJ (2000). Identification of Ubiquilin, a Novel Presenilin Interactor That Increases Presenilin Protein Accumulation. J Cell Biol 151, 847–862.

Mann, JR et al. (2019). RNA Binding Antagonizes Neurotoxic Phase Transitions of TDP-43. Neuron 102, 321–338.e8.

Marzahn, MR et al. (2016). Higher-order oligomerization promotes localization of SPOP to liquid nuclear speckles. EMBO J 35, 1254–1275.

Molliex, A, Temirov, J, Lee, J, Coughlin, M, Kanagaraj, AP, Kim, HJ, Mittag, T, and Taylor, JP (2015). Phase Separation by Low Complexity Domains Promotes Stress Granule Assembly and Drives Pathological Fibrillization. Cell 163, 123–133.

Monahan, Z et al. (2017). Phosphorylation of the FUS low-complexity domain disrupts phase separation, aggregation, and toxicity. EMBO J 36, 2951–2967.

Morelli, FF et al. (2017). Aberrant Compartment Formation by HSPB2 Mislocalizes Lamin A and Compromises Nuclear Integrity and Function. Cell Rep 20, 2100–2115.

Osaka, M, Ito, D, and Suzuki, N (2016). Disturbance of proteasomal and autophagic protein degradation pathways by amyotrophic lateral sclerosis-linked mutations in ubiquilin 2. Biochem Biophys Res Commun 472, 324–331.

Patel, A et al. (2015). A Liquid-to-Solid Phase Transition of the ALS Protein FUS Accelerated by Disease Mutation. Cell 162, 1066–1077.

Picher-Martel, V, Dutta, K, Phaneuf, D, Sobue, G, and Julien, J-P (2015). Ubiquilin-2 drives NF-κB activity and cytosolic TDP-43 aggregation in neuronal cells. Mol Brain 8, 71.

Schindelin, J et al. (2012). Fiji: an open-source platform for biological-image analysis. Nat Methods 9, 676–682.

Sharkey, LM et al. (2018). Mutant UBQLN2 promotes toxicity by modulating intrinsic self-assembly. Proc Natl Acad Sci 115, E10495–E10504.

Taylor, JP, Brown Jr, RH, and Cleveland, DW (2016). Decoding ALS: from genes to mechanism. Nature 539, 197–206.

Teyssou, E et al. (2017). Novel UBQLN2 mutations linked to Amyotrophic Lateral Sclerosis and atypical Hereditary Spastic Paraplegia phenotype through defective HSP70-mediated proteolysis. Neurobiol Aging 58, 239.e11–239.e20.

Whiteley, AM et al. (2020). Global proteomics of Ubqln2-based murine models of ALS. BioRxiv, 2020.02.22.956524.

Wu, JJ et al. (2020). ALS/FTD mutations in UBQLN2 impede autophagy by reducing autophagosome acidification through loss of function. Proc Natl Acad Sci.

Wu, Q, Liu, M, Huang, C, Liu, X, Huang, B, Li, N, Zhou, H, and Xia, X-G (2015). Pathogenic Ubqln2 gains toxic properties to induce neuron death. Acta Neuropathol (Berl) 129, 417–428.

Xia, Y, Yan, LH, Huang, B, Liu, M, Liu, X, and Huang, C (2014). Pathogenic mutation of UBQLN2 impairs its interaction with UBXD8 and disrupts endoplasmic reticulum-associated protein degradation. J Neurochem 129, 99–106.

Yang, P et al. (2020). G3BP1 Is a Tunable Switch that Triggers Phase Separation to Assemble Stress Granules. Cell 181, 325–345.e28.

