## Supporting Information for "ALS-linked mutations impair UBQLN2 stress-induced biomolecular condensate assembly in cells"

### Supplementary Materials

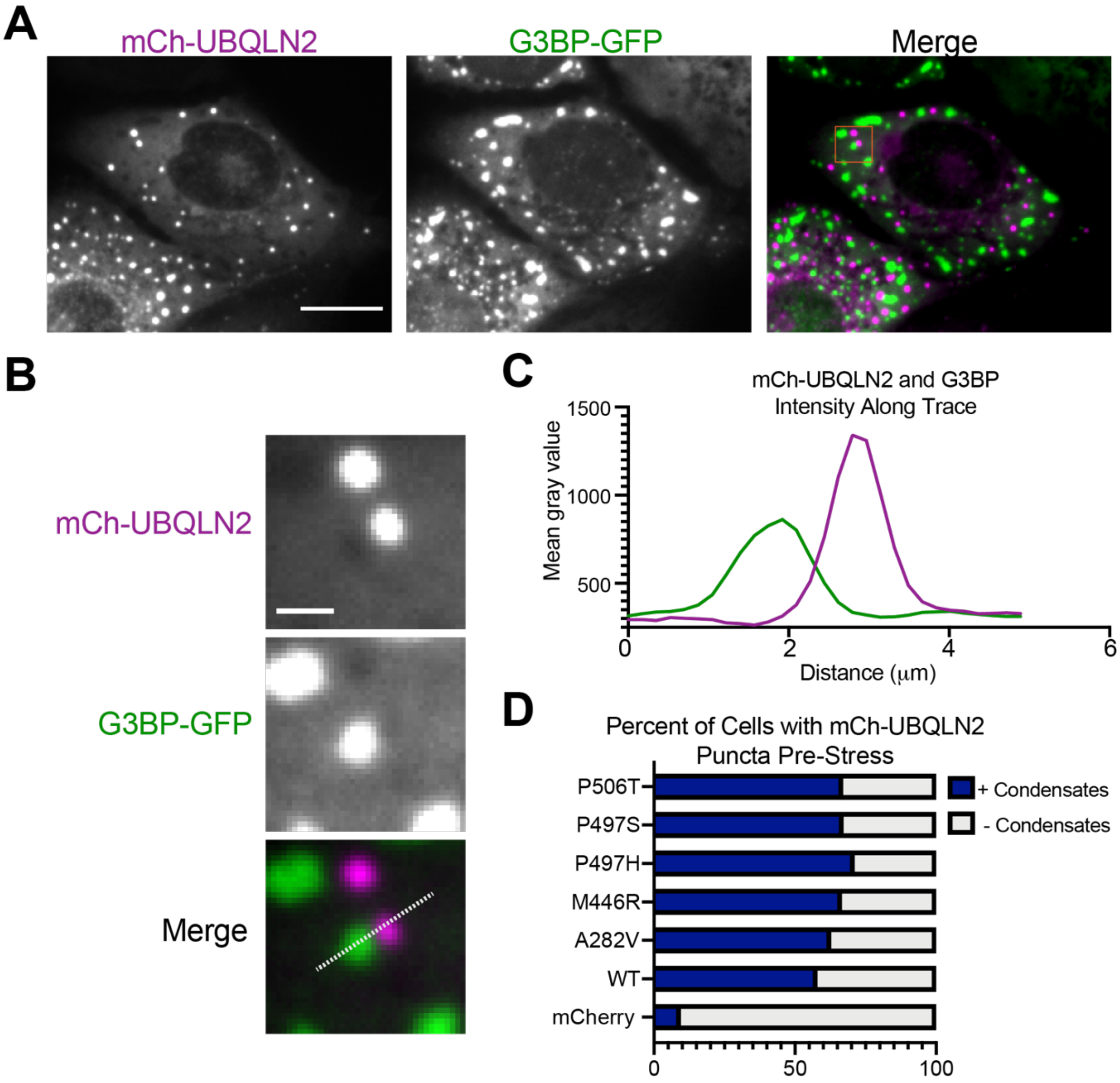


**Figure S1. mCh-UBQLN2 bodies are distinct from stress granules.** (A) mCh-UBQLN2 and stress granule marker G3BP-GFP 60 minutes following the application of 0.5mM NaAsO2. Scale bar 15 μm. (B) mCh-UBQLN2 and G3BP-GFP images for area enclosed in an orange box on the merged image in (A). Scale bar 2 μm. (C) Mean intensity of mCh-UBQLN2 (magenta) and G3BP-GFP (green) measured along the white line in (B). (D) Average percent of cells transfected with each variant of mCh-UBQLN2 or a mCherry control that had puncta prior to the application of arsenite stress.

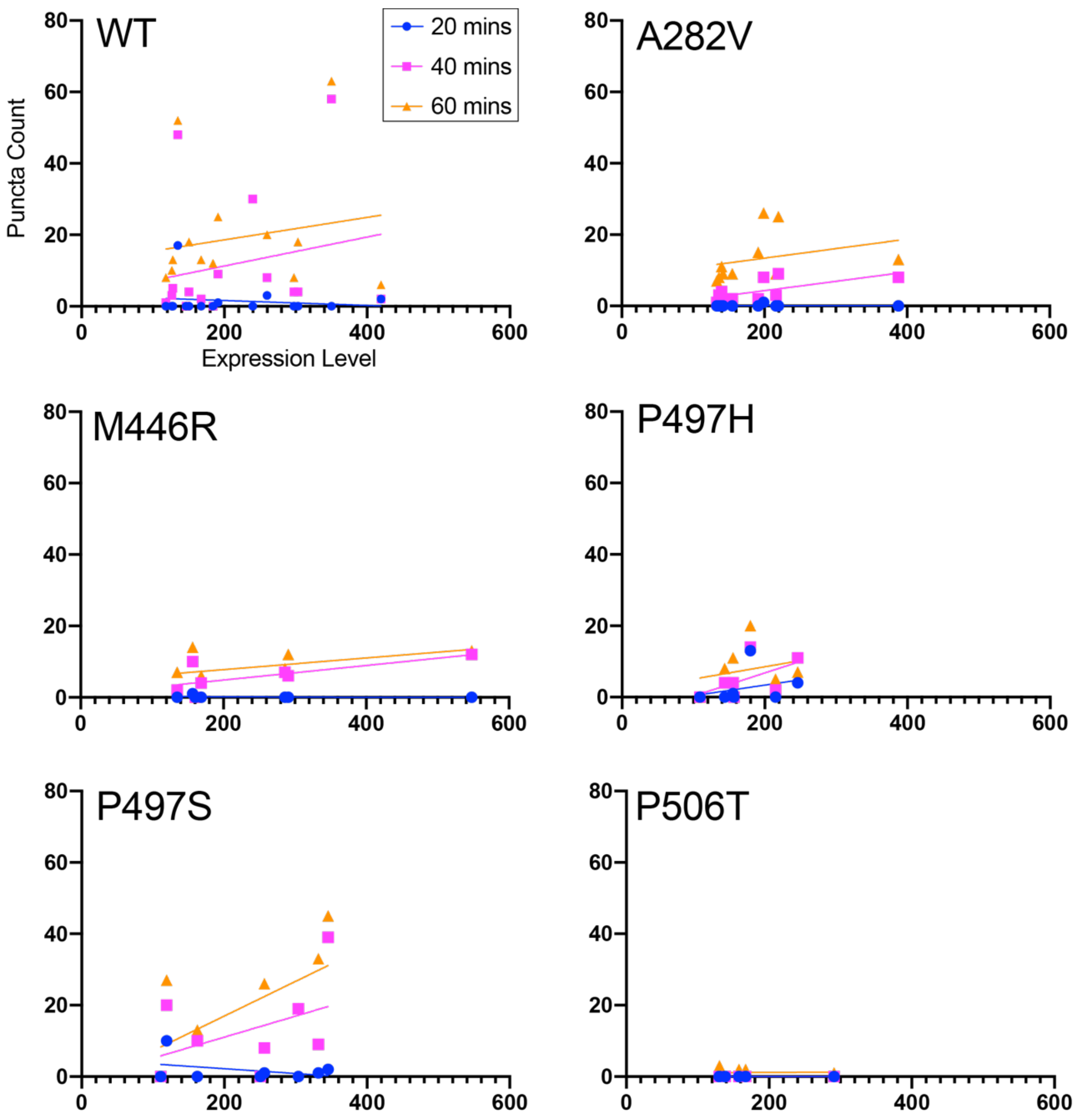


**Figure S2. mCh-UBQLN2 expression levels do not correlate with number of UBQLN2 bodies that form in response to arsenite stress.** Number of stress-induced UBQLN2 bodies at 20, 40 and 60 minutes post-$NaAsO_2$ addition as a function of mean intensity of mCh-UBQLN2 in each cell. WT n=15, $R^2$ (Pearson's correlation coefficient) values of 0.021 (20 min), 0.042 (40 min), and 0.030 (60 min); A282V n=10, $R^2$ values of 0.000 (20 min), 0.426 (40 min), and 0.087 (60 min); M446R n=7, $R^2$ values of 0.078 (20 min), 0.488 (40 min) and 0.237 (60 min); P497H n=7, $R^2$ values of 0.080 (20 min), 0.299 (40 min) and 0.057 (60 min); P497S n=8, $R^2$ values of 0.135 (20 min), 0.186 (40 min) and 0.335 (60 min); P506T n=7, $R^2$ value of 0.001 (60 min, the only time point where P506T mCh-UBQLN2 formed any puncta).

### Supplementary Movies

**Movie S1. P497H mCh-UBQLN2 condensate formation.** P497H mCh-UBQLN2 stress-induced puncta have liquid-like properties as marked puncta coalesce over time and return to a round shape. Scale bar 15 μm. Movie is approximately 70 minutes of real time (4 frames/sec).

**Movie S2. P497S mCh-UBQLN2 condensate formation.** P497S mCh-UBQLN2 stress-induced puncta have liquid-like properties as marked puncta coalesce over time and return to a round shape. Scale bar 15 μm. Movie is approximately 70 minutes of real time (4 frames/sec).
